# Resting EEG power spectra across middle to late life: Associations with age, cognition, APOE-ε4 carriage and cardiometabolic burden

**DOI:** 10.1101/2022.09.01.506258

**Authors:** Ashleigh E. Smith, Anson Chau, Danielle Greaves, Hannah A.D. Keage, Daniel Feuerriegel

## Abstract

We investigated how resting EEG measures are associated with risk factors for late-life cognitive impairment and dementia, including age, Apolipoprotein E ε4 (APOE-ε4) carriage and cardiometabolic burden. Resting EEG was recorded from 86 adults (50-80 years of age). Participants additionally completed the Addenbrooke’s Cognitive Examination (ACE) III and had blood drawn to assess APOE-ε4 carriage status and cardiometabolic burden. EEG power spectra were decomposed into sources of periodic and aperiodic activity to derive measures of aperiodic component slope and alpha (7-14 Hz) and beta (15-30 Hz) peak power and peak frequency. Alpha and beta peak power measures were corrected for aperiodic activity. The aperiodic component slope was correlated with ACE-III scores but not age. Alpha peak frequency decreased with age. Individuals with higher cardiometabolic burden had lower alpha peak frequencies and lower beta peak power. APOE-ε4 carriers had lower beta peak frequencies. Our findings suggest that the slope of the aperiodic component of resting EEG power spectra is more closely associated with measures of cognitive performance rather than chronological age in older adults.

## 1. Introduction

A combination of non-modifiable and modifiable factors is associated with our risk of developing late life cognitive impairment and dementia. Increased age and carriage of the Apolipoprotein E ε4 (APOE-ε4) gene are the largest non-modifiable risk factors (Raber et al., 2004; Boyle et al., 2019). In addition, it has been estimated that 40% of dementia risk in people over 65 years is attributable to modifiable lifestyle risk factors including Type II diabetes, obesity, physical inactivity, hypertension and high cholesterol (Norton et al., 2014; Livingston et al., 2020). In this study, we investigated how non-modifiable factors (age and APOE-ε4 carriage) and modifiable factors (cardiometabolic burden) associated with brain function and cognitive performance in adults in mid- to late-life.

Electroencephalography (EEG) is a non-invasive and widely available tool for measuring dynamic patterns of brain activity (primarily postsynaptic potentials from pyramidal neurons) using electrodes placed on the scalp (Buzsáki and Draguhn, 2004). EEG recorded at rest is often converted into frequency domain representations using Fourier transforms to assess individual or group differences in spectral power profiles. In such analyses, researchers have typically used measures of summed power within fixed frequency bands (e.g., delta, theta, alpha, beta, or gamma). Using this approach, they have reported decreases in power across the delta (<4Hz) and alpha (7-13Hz) bands with increased age, as well as a slowing of alpha peak frequency (e.g., Celesia, 1986; Klimesch, 1997; Babiloni et al., 2006b; Vlahou et al., 2015; Sghirripa et al., 2021; reviewed in Scally et al., 2018). Age-related increases in theta band power across mid- to late-life have also been reported, however these changes have been attributed to brain activity typically associated with the alpha-band occurring at slower frequencies in older adults (reviewed in Klimesch, 1999; Finley et al., 2022).

In previous work, researchers have suggested that certain factors (which become more prevalent with increased age) are also associated with changes to resting EEG spectra. These factors include the presence of mild cognitive impairment (MCI) and dementia (Babiloni et al., 2013) and severe cardiovascular complications such as congestive heart failure (Vecchio et al., 2012). Compared to age-matched controls, people with congestive heart failure showed increased delta and decreased alpha power (Vecchio et al., 2012). These differences, as well as a slowing of alpha peak frequency, were seen in individuals with Alzheimer’s Disease, and to a lesser extent for MCI (Babiloni et al., 2013; Meghdadi et al., 2021; Vecchio et al., 2013). We have also demonstrated that the P3b event-related potential (ERP) component recorded during an n-back memory task is reduced in older adults with increased cardiometabolic burden (Keage et al., 2020). Taken together, this evidence suggests that there are partially overlapping EEG correlates of increased age, cognitive decline and cardiometabolic burden.

One important shortcoming of the analysis approaches described above is that they often consider frequency band power measures as directly reflecting periodic (or oscillatory) patterns of neural activity, and do not consider the contribution of aperiodic EEG activity (sometimes labeled ‘background noise’) to Fourier power spectra (Donoghue et al., 2020a; Schaworonkow and Voytek, 2021). Aperiodic EEG activity exhibits a spectral profile that decreases in power with increasing frequency, which can be approximated using exponentially decreasing functions. Typical frequency band power measures conflate contributions of aperiodic and periodic activity. In addition, the magnitude of the aperiodic component — contributing to band power measures across a wide range of frequencies, and typically indexed by an offset parameter when modeling power spectra — can vary across individuals in systematic ways. For example, the magnitude of aperiodic activity is larger in younger compared to older adults (e.g., Donoghue et al., 2020a; Merkin et al., 2021), meaning that frequency band power measures are also likely to be larger in younger adult groups. The shape of the aperiodic component (i.e. the rate of decrease in power with increasing frequency, here termed the aperiodic slope) also appears to be closely linked to band power ratio measures used in resting EEG research (Donoghue et al., 2020b). For example, Finley et al. (2022) reported that the ratio of power in the theta compared to the beta band was lower in older adults, but this association was largely due to changes in the shape of the aperiodic component. This example underlines the importance of separately modeling aperiodic and periodic components of Fourier power spectra (for further discussion see Donoghue et al., 2020a).

Recent studies that have isolated aperiodic and periodic contributions do not report such widespread effects of age on periodic EEG measures when aperiodic activity is accounted for (Voytek et al., 2015; Merkin et al., 2021). Instead, the slope of the spectral profile of the aperiodic component (the aperiodic slope) has been found to systematically vary with age, becoming steeper throughout childhood (He et al., 2019) and flatter during adolescence (McSweeney et al., 2021) and with increasing age during adulthood (Voytek et al., 2015; Dave et al, 2018; Leenders et al., 2019; McNair et al., 2019; Kosciessa et al., 2020; Tran et al., 2020; Merkin et al., 2021). Here, we note that the aperiodic slope is distinct from the overall magnitude of aperiodic activity measured using an offset parameter. The latter findings have been proposed to support the neural noise hypothesis of aging (Cremer & Zeef, 1987; Voytek & Knight, 2015), which suggests that the balance between excitatory and inhibitory neural activity changes across adulthood, leading to a flatter slope of the aperiodic component of the EEG power spectrum (Hong and Rebec, 2012; Gao et al., 2017). Under this hypothesis, the flattening of the aperiodic slope is also associated with a reduction of the signal to noise ratio of neuronal communication, resulting in a decline in cognitive performance (Voytek & Knight, 2015; Voytek et al., 2015; Dave et al., 2018).

It remains an open question whether these findings are specifically due to changes with increased chronological age, or other changes which co-occur with aging across early- to late-adulthood. Almost all of the studies listed above compared younger and older groups of markedly different age ranges (e.g., 20-30 year-olds compared to 60-70 year-olds in Voytek et al., 2015). Studies treating age as a continuous variable have recruited very small samples (n=15 in Voytek et al., 2015, n=19 in Waschke et al., 2017), making it difficult to assess the functional form of age-aperiodic slope associations (but see Finley et al., 2022 for an approximately linear association in a large sample of 36-84 year-olds). Importantly, flatter aperiodic slopes have also been reported in individuals with worse cognitive performance (Voytek et al., 2015; Ouyang et al., 2020), and cognitive performance in many tasks steadily declines with increasing age.

Consequently, it is difficult to assess whether flatter aperiodic slopes are specifically associated with increasing chronological age, or other factors that co-occur with aging and likely to affect brain function, such as changes in cardiometabolic burden or neurovascular coupling (e.g., Han et al., 2020). Disentangling these different effects is useful for refining the neural noise hypothesis of aging, which describes both increased age and cognitive performance as associated with the shape of the aperiodic component of neural power spectra.

In the current study, we recruited adults between 50 and 80 years of age and used analysis techniques to disentangle aperiodic and periodic contributions to resting EEG power spectra. We tested for associations between chronological age, cognitive performance, and the slope of the aperiodic component. We also tested for previously reported associations between age, cognitive performance, and alpha peak frequency (e.g., Klimesch, 1999; Grandy et al., 2013; Scally et al., 2018; Sghirripa et al., 2021; Merkin et al., 2021). In line with published work, we hypothesized that, with increased age, the slope of the aperiodic component would be flatter and alpha peak frequency would be lower. We also conducted a broad search for resting EEG correlates of APOE-ε4 carriage and cardiometabolic burden while accounting for effects of age. These analyses were exploratory in nature to identify candidate associations that could be replicated in future work.

## 2. Methods

### 2.1. Participants

Participants were deemed eligible if they were aged between 50 and 80 years and had no known history of stroke, clinical dementia or current psychiatric disorder diagnosis, blindness or vision problems not corrected by glasses/contact lenses, a previous period of unconsciousness that lasted greater than 5 minutes, or any known intellectual disabilities. To gain a broad sample distribution of cardiometabolic disease risk burden, participants were selectively recruited based on their self-reported cardiometabolic burden using the online Framingham risk calculator (Harrison et al., 2017; Wolf et al., 1991). Within each decade of 50-59, 60-69, and 70-79 years of age we intended to recruit 30 participants (15 with self-reported low cardiovascular disease risk and 15 self-reported high risk).

Participants were recruited as part of a larger study investigating the neurophysiological and lifestyle changes across mid to late life. Other results using task-based EEG and dietary measures are published elsewhere (Keage et al., 2020; Wade et al., 2020). The study was approved by the University of South Australia Human Ethics Committee (0000034635).

### 2.2. Procedure

All participants attended two three-hour appointments separated by approximately 8-10 days. In session one, informed consent, general health, cognition, fasted blood tests (minimum 8-hour fast), anthropometric assessments, blood pressure (arterial compliance measurement), and dietary assessments were conducted. In session two, EEG data was recorded both during rest and then during a cognitive task, however only resting EEG data are presented here.

### 2.3. Measures

#### 2.3.1. EEG recording and processing

Two minutes of eyes open and two minutes of eyes closed resting EEG data were collected. The order of eyes open/closed conditions was counterbalanced across participants. We recorded EEG from 25 active electrodes (Fp1, Fp2, AFpz, Fz, F3, F7, T7, C3, Cz, Pz, P3, P7, PO7, PO3, O1, Oz, O2, PO4, PO8, P8, P4, C4, T8, F8, F4) using a Biosemi Active Two system (Biosemi, the Netherlands). Recordings were grounded using common mode sense and driven right leg electrodes (http://www.biosemi.com/faq/cms&drl.htm). EEG was sampled at 1024Hz (DC-coupled with an anti-aliasing filter, -3dB at 204Hz). Electrode offsets were kept within ±50μV. We processed EEG data using EEGLab V.13.4.4b (Delorme and Makeig, 2004) running in MATLAB (The Mathworks).

EEG data were re-referenced offline to a nose reference. Data were high-pass filtered at 1 Hz (EEGLab Basic FIR Filter New, zero-phase, finite impulse response, -6 dB cutoff frequency 0.5 Hz, transition bandwidth 1 Hz) and then low-pass filtered at 30 Hz (EEGLab Basic Finite Impulse Response Filter New, zero-phase, -6 dB cutoff frequency 33.75 Hz, transition band width 7.5 Hz). Excessively noisy channels were identified by visual inspection and were not included as input data for the independent components analysis (ICA). Data were then segmented into epochs spanning 0 to 120 seconds relative to the start of each eyes open and eyes closed condition. ICA was performed on the resulting data (RunICA extended algorithm, Jung et al., 2000). Independent components associated with eyeblinks and saccades were identified and removed according to guidelines in Chaumon et al. (2015). Excessively noisy channels were then interpolated using the cleaned data (using spherical spline interpolation).

Fourier power spectra were calculated for each channel using Welch’s method (2-second Hann windowed epochs, 50% overlap, 0.5 Hz frequency resolution) for eyes open and eyes closed conditions separately. During this step, epochs with amplitudes exceeding ±150μV were removed prior to averaging across epochs. We then used the fitting oscillations & one over f (FOOOF) algorithm (Donoghue et al., 2020) to parameterize the resulting power spectra into aperiodic and periodic components, using the frequency range of 2-30 Hz (no knee parameter, periodic component peak width limits of 1Hz and 8Hz, minimum peak height = 0.05, maximum of 6 peaks identified per power spectrum). Power spectra were modeled for each electrode separately, and for the eyes open and closed conditions separately. We extracted the aperiodic exponent (where larger exponent values indicate steeper slopes) and the aperiodic offset (where larger values indicate higher overall power across the Fourier power spectra), and also attempted to extract peak power and peak frequency values within the delta (2-3 Hz), theta (4-7 Hz), alpha (7-14 Hz) and beta (15-30 Hz) frequency bands. We used a wide frequency range for the alpha band compared to some existing studies on younger adults because lower peak frequency values have been reported in older adults (e.g., Ishii et al., 2017; Mizukami & Katada, 2018).

Importantly, peak power values were corrected for the contribution of aperiodic activity to the Fourier power spectra. The algorithm could only reliably identify peaks across participants in the alpha and beta bands, and clear peaks in other bands were not visible in the group-averaged power spectra. Consequently, peak power and peak frequency measures were not extracted for the delta and theta bands (for similar results from resting EEG recordings in older adults see Merkin et al., 2021). Estimates of the aperiodic exponent and alpha and beta peak power values and peak frequencies, calculated for each electrode separately, were then averaged across electrodes within pre-defined regions of interest (ROIs). Different ROIs (i.e., different sets of electrodes) were used for each measure derived from the parameterised power spectra. This resulted in one value of each ROI-averaged measure per eyes open/closed condition per participant. For alpha peak frequencies and power values we used a parietal ROI consisting of channels Pz, P3, P4, PO3 and PO4. This ROI included parietal and occipito-parietal electrodes because alpha band power is known to show a focal distribution over these channels (Klimesch, 1999; Merkin et al., 2021). For the aperiodic exponent, aperiodic offset and beta band peak frequencies and power values (which are not known to have a clearly defined focal distribution on the scalp) we included a broad array of electrodes (excluding channels on the edge of the cap) consisting of Fz, F3, F4, Cz, C3, C4, Pz, P3, P4, PO3 and PO4. Here, we label this the broad ROI as it includes electrodes spanning frontal, central and parietal sites.

To enable comparisons with prior work using conventional band power measures we also calculated the average power within the delta, theta, alpha and beta bands. Methodological details and results are provided in the Supplementary Material.

Estimates from eyes open and eyes closed conditions comprised separate data points for each participant, and both were included in the resulting statistical analyses. For Pearson and Spearman correlation analyses we first averaged measures across eyes open and closed conditions within each participant. For mixed-effects modeling we included measures from both eyes open and closed conditions, and treated eyes open/closed condition as a within-subject factor. We included both conditions to increase data quantity and the precision of estimates, and because we did not have a clear a priori reason (based on existing theory or previous work) to expect different patterns of associations with age or other variables across these conditions.

#### 2.3.2. Cardiometabolic burden and APOE analysis

Blood glucose, waist to hip ratio (obesity), total blood cholesterol, and blood pressure were used to objectively characterize cardiometabolic disease risk burden. Notably, we measured total blood cholesterol rather than high-density lipoprotein (HDL) or low-density lipoprotein (LDL), as it is this measure that demonstrates associations with incident cognitive impairments in late life such as MCI and dementia (Anstey et al., 2017). Participants were classified as positive for each cardiometabolic factor based on the following cut-offs: blood glucose ≥6.5 mmol/L (Carson et al., 2010; Howe-Davies et al., 1980); waist to hip ratio ≥.95 for males or ≥.90 for females (Gill et al., 2003; WHO, 2011); total blood cholesterol ≥5.5 mmol/L (Solomon et al., 2009); and if either diastolic blood pressure (BP) was ≥90 mmHg or systolic blood pressure was ≥140 mmHg (Muntner et al., 2018; Whelton et al., 2018). Although there are various cut-offs reported across the previous literature, we used the most frequently reported.

Waist to hip ratios were obtained by research assistants trained in the standard protocols used by the International Society for the Advancement of Kinanthropometry (Stewart et al., 2011). Using a Luftkin executive thin line 2mm metal tape measure, waist measures were taken at the point of visible narrowing between the 10^th^ rib and the crest of the ilium during normal expiration. In the event there was no narrowing the measurement was taken at the mid-point between the lower costal (10^th^ rib) border and iliac crest. Hip measures were taken at the greatest point of posterior protuberance of the buttocks. Two separate measures were taken and if the measures differed by >20% a third measure was taken.

Prior to blood pressure measurements participants rested horizontally for a minimum of five minutes in a dark room. Blood pressure was measured non-invasively in conjunction with arterial compliance using a cardiovascular profiler (HDI cardiovascular profiler CR 2000, Hypertension Diagnostics, Minnesota, United States). The blood pressure cuff was fitted over the left brachial artery. Three readings were performed at 5-minute intervals and the average reading was calculated. If readings differed >20%, an additional reading was completed.

Whole blood was collected via venepuncture into 2 × 9mL ethylenediaminetraacetic acid (EDTA) (18mg) anticoagulant and 1 × 4mL sodium fluoride Vacuette tubes (grenier bio-one, Kremsmünster, Austria). Collected samples were centrifuged at 4000 rpm for 10 minutes, aliquoted into Eppendorf tubes and frozen initially at -20°C for up to one week before being transferred to and stored at -80°C until analysis.

Cholesterol, triglycerides, HDL and glucose (from serum sample) were analyzed in duplicate with a commercial assay kit (including quality controls and calibrators) using the KONELAB 20XTi (ThermoFisher, Massachusetts, United States). Deoayribonucleic acid (DNA) was extracted from stored buffy coat using the Blood Mini DNA kit (Qiagen, Valencia, CA). A TaqMan^®^ single nucleotide polymorphism (SNP) genotyping assay (Applied Biosystems, California, USA) was used to determine APOE SNPs rs429358 (Cys112Arg) and rs7412 (Arg158Cys). This involved setting up a 96 well plate with 1ng/μL of DNA, 1 x TaqMan® Universal polymerase chain reaction (PCR) Master Mix and 1 x genotype assay mix (containing the fluorescently labeled primers Vic and Fam for allele identification) (Applied Biosystems, California, United States). PCR conditions included an initial holding stage at 95°C for 10 minutes; 40 cycles of 15 seconds at 95°C followed by 60°C for 1 minute; holding stage of 60°C for 1 minute (Applied Biosystems 7500 Fast Real-Time PCR System, California, United States). Non template controls were performed in triplicate for each SNP and plate. Participants were classified as APOE-ε4 carriers if they were heterozygous *or* homozygous (i.e., carrying one or both APOE-ε4 alleles) as per Ghebranious et al. (2005). Participants who did not carry an APOE-ε4 allele were classified as noncarriers (Lim et al., 2015).

#### 2.3.3. Cognitive performance

The Addenbrooke’s Cognitive Examination III (ACE-III) was administered to assess global cognition and as a broad screening for cognitive impairment and dementia (Hsieh et al., 2013). The ACE-III consists of five subscales assessing attention/orientation (18 points), memory (26 points), fluency (14 points), language (26 points), and visuospatial abilities (16 points); therefore, possible score range of 0 to 100, with a higher score reflecting higher cognitive function. The test takes approximately 15-20 minutes to administer, and has demonstrated (along with its predecessor, the ACE-Revised) high sensitivity (1.0) and specificity (0.96) and very good internal reliability, with an alpha coefficient of 0.8 (Hsieh et al., 2013; Mioshi et al., 2006).

### 2.4. Statistical Analyses

Statistical analyses were conducted with JASP v0.14.1 for correlation analyses, and STATA v15.1 IC for mixed effects modeling. EEG peak power variables were log transformed.

We tested for bivariate associations between aperiodic exponent values and age and ACE-III scores, between alpha peak frequencies and age and ACE-III scores, and between age and ACE-III scores using Pearson correlations. In cases where bivariate normality assumptions were violated, results from Spearman correlations are also reported. Exponent and alpha peak frequency values were averaged across eyes open and eyes closed conditions for each participant prior to correlation analyses. Bayes Factors in favor of the alternative hypothesis were also calculated using JASP (stretched beta prior distribution, width = 1.0). Please note that these analyses do not include covariates and are simple pairwise correlations, consistent with those used in existing work (e.g., Voytek et al., 2015; Finley et al., 2022). In a separate set of exploratory analyses we also used Pearson correlations to test for associations between aperiodic offsets (using measures averaged over channels within the broad ROI), age and ACE scores.

To identify effects of APOE-ε4 carriage and cardiometabolic burden while accounting for effects of age, mixed effects modeling with maximum likelihood estimation, with participant identity set as both a random intercept and slope, was conducted for each EEG measure of interest: exponent, alpha peak frequency, alpha peak power, beta peak frequency and beta peak power. Predictor variables across all models included: cardiometabolic burden (0=none, 1=one, 2=two or more factors above cut-off), condition (1=eyes open, 2=eyes closed) and the presence of the APOE-ε4 allele (1=no E4 allele, 2=presence of at least one E4 allele). Cohen’s *f*^*2*^ was our measure of effect size, with *f*^*2*^ values > .02, >.15, and >.35 representing small, medium, and large effect sizes, respectively. We did not employ a multiple testing correction due to the exploratory nature of our approach. We additionally fitted mixed-effects models using subgroups of participants that scored below the 92 cutoff for MCI in the ACE-III (results displayed in Supplementary Table 1) and above the cutoff (Supplementary Table 2). However, we note that scores on the ACE-III do not indicate an MCI diagnosis, and no participants had a formal diagnosis of MCI.

### 2.5. Data Availability

Code and data used in the analyses reported in this paper will be available at https://osf.io/jasrb/ at the time of publication.

## 3. Results

We were unable to record EEG for four participants and obtained only one usable epoch of data from one participant, resulting in five excluded participants in total. Therefore, the total number of participants included in these analyses was 86 (51 female, age range 50-80; mean age 65.5 ± 7.7 years). These exclusion judgments were made prior to data analysis. Over half (55%) of participants scored below the 92 ACE-III MCI cut-off (Pendlebury et al., 2012; North et al. 2021). However, please note that no participants had a formal MCI diagnosis, and that ACE-III scores can be influenced by factors such as one’s level of education (and we did not record these data). ACE-III scores ranged between 72 and 99, with a mean of 91.3 (SD=5.3).

### 3.1. Distribution of Cardiometabolic Burden and APOE-ε4 Carriage

Table 1 displays the distribution of cardiometabolic burden relative to each of the four cardiometabolic factor cut-offs, and the summative variable (number of factors above the cut-off: 0, 1 or 2 or more). Participants were sampled so that approximately equivalent numbers of participants were included with 0, 1 and 2 or more risk factors. APOE genotype was determined for 80 out of 86 participants, with n=22 (27.5%) displaying the presence of at least one ε4 allele.

**Table 1.** Distribution of cardiometabolic burden across the sample.

| | High blood<br>glucose<br>($\geq 6.5$<br>mmol/L) | High<br>waist:hip<br>($\geq .95$ men/<br>$\geq .90$<br>women) | High total<br>blood<br>cholesterol<br>($\geq 5.5$<br>mmol/L) | Hypertension<br>(diastolic<br>$\geq 90$/<br>systolic $\geq 140$<br>mmHg) | Summative<br>cardiometabolic burden | | |
| --- | --- | --- | --- | --- | --- | --- | --- |
| | | | | | 0 | 1 | $\geq 2$ |
| Female<br>($n=51$) | 9<br>(18%) | 9<br>(18%) | 20<br>(39%) | 13<br>(25%) | 16<br>(31%) | 22<br>(43%) | 13<br>(25%) |
| Male<br>($n=35$) | 14<br>(40%) | 20<br>(57%) | 8<br>(23%) | 12<br>(35%) | 5<br>(14%) | 12<br>(34%) | 18<br>(51%) |
| Total<br>($n=86$) | 23<br>(27%) | 29<br>(34%) | 28<br>(33%) | 25<br>(29%) | 21<br>(24%) | 34<br>(40%) | 31<br>(36%) |

### 3.2. Characteristics of EEG Fourier Power Spectra for Eyes Open and Eyes Closed Conditions

Fourier power spectra were typical of those in eyes open and eyes closed resting conditions (Figure 1A). As revealed by mixed effects modeling (results in Table 2), in the eyes closed condition there were larger exponents (i.e., steeper aperiodic component slopes), larger peak power values in the alpha and beta bands, and lower peak frequencies in the alpha and beta bands, as compared to the eyes open condition. Please note that alpha and beta peak power values are corrected for the contribution of aperiodic activity to the Fourier power spectra. Figure 1B shows the topographical distributions of group-averaged estimates of the exponent, as well as alpha and beta peak power and peak frequency values. Values of the exponent were similar across a broad region of the scalp, consistent with prior observations (e.g., Tran et al., 2020; Merkin et al., 2021). By contrast, alpha band peak power and frequency values were largest over parietal channels. Beta band peak power estimates were largest around parietal electrodes, however peak frequencies in the beta band tended to be lowest in this parietal region as compared to more frontal electrodes.

**Figure 1.**
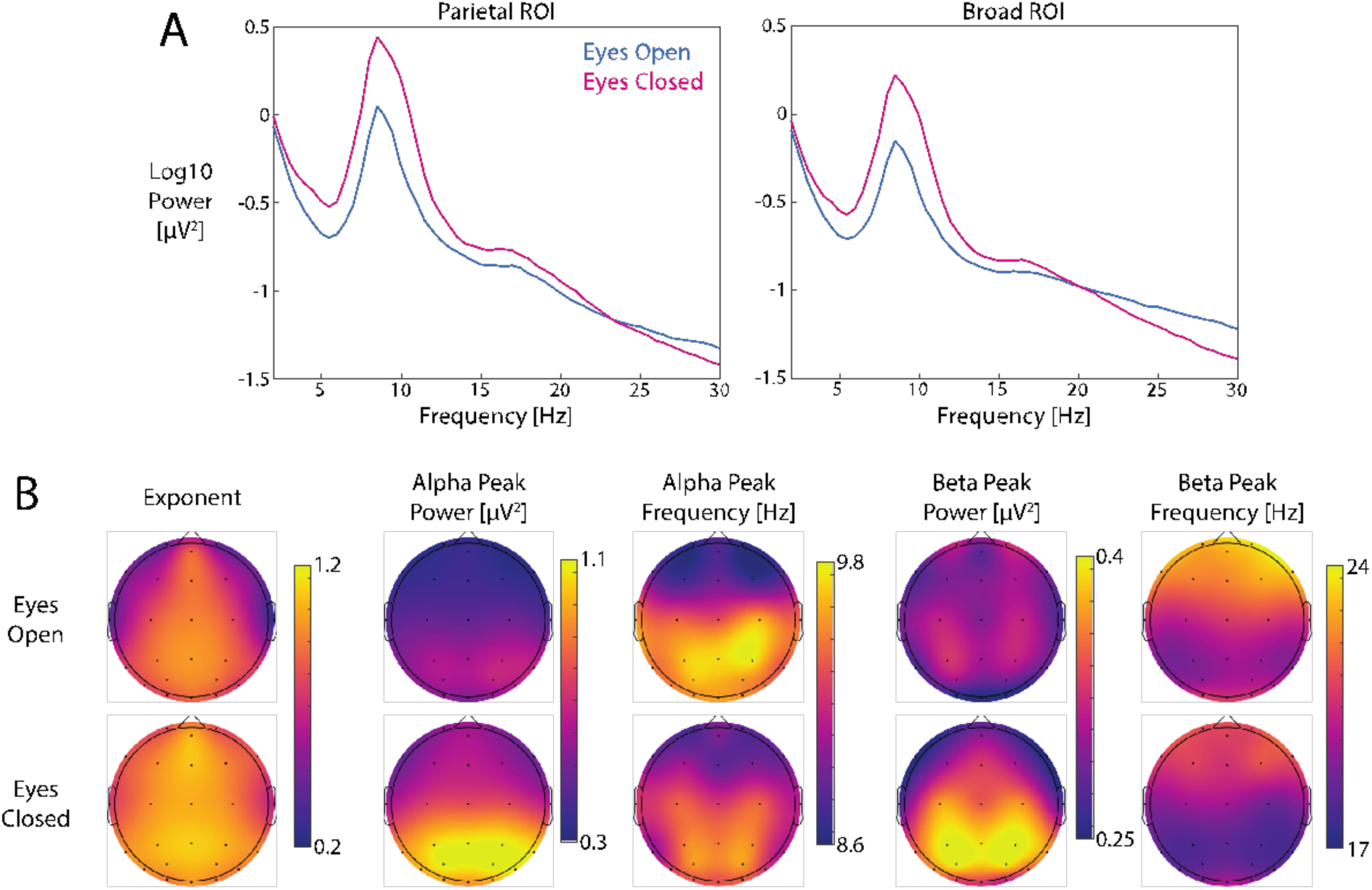
Characteristics of the EEG power spectra in the sample. A.) Fourier power spectra for eyes open and eyes closed conditions, averaged over electrodes included in the parietal ROI (left plot, used to measure alpha peak power and peak frequency) and the broad ROI (right plot, used to measure the exponent and beta peak power and peak frequency). The parietal ROI includes electrodes Pz, P3, P4, PO3 and PO4. The broad ROI includes electrodes Fz, F3, F4, Cz, C3, C4, Pz, P3, P4, PO3 and PO4. B.) Scalp maps of group-averaged estimates of the exponent, alpha peak power, alpha peak frequency, beta peak power and beta peak frequency, for eyes open (top row) and eyes closed (bottom row) conditions. Please note that alpha and beta peak power values are corrected for contributions of aperiodic activity.

**Table 2.** Results from mixed models, with EEG indices as outcomes, and predictors including condition (eyes open/eyes closed), age, cardiometabolic burden and APOE-ε4 status. Statistically significant (<.05) p-values are shown in bold text.

|  | beta | SE | P | 95% CI |  | f<sup>2</sup> |
| --- | --- | --- | --- | --- | --- | --- |
| Exponent |  |  |  |  |  |  |
| EO (1) v EC (2) | 0.154 | 0.029 | <.001 | 0.097 | 0.211 | 0.350 |
| Age in years | -0.005 | 0.004 | .196 | -0.012 | 0.002 | <0.001 |
| Cardiometabolic burden | -0.268 | 0.036 | .458 | -0.098 | 0.044 | <0.001 |
| APOE-ε4 | 0.034 | 0.059 | .565 | -0.082 | 0.151 | <0.001 |
| Alpha peak frequency |  |  |  |  |  |  |
| EO (1) v EC (2) | -0.249 | 0.123 | .043 | -0.491 | -0.008 | 0.050 |
| Age in years | -0.036 | 0.013 | .007 | -0.062 | -0.010 | <0.001 |
| Cardiometabolic burden | -0.277 | 0.136 | .042 | -0.544 | -0.010 | <0.001 |
| APOE-ε4 | -0.262 | 0.224 | .241 | -0.700 | 0.176 | 0.030 |
| Alpha peak power (log) |  |  |  |  |  |  |
| EO (1) v EC (2) | 0.668 | 0.066 | <.001 | 0.538 | 0.798 | 1.279 |
| Age in years | 0.009 | 0.008 | .281 | -0.007 | 0.025 | <0.001 |
| Cardiometabolic burden | -0.112 | 0.082 | .170 | -0.272 | 0.048 | 0.001 |
| APOE-ε4 | -0.047 | 0.133 | .727 | -0.307 | 0.214 | <0.001 |
| Beta peak frequency |  |  |  |  |  |  |
| EO (1) v EC (2) | -1.318 | 0.254 | <.001 | -1.816 | -0.820 | 0.335 |
| Age in years | 0.053 | 0.037 | .149 | -0.019 | 0.124 | 0.002 |
| Cardiometabolic burden | 0.709 | 0.367 | .054 | -0.012 | 1.429 | <0.001 |
| APOE-ε4 | -1.250 | 0.595 | .036 | -2.417 | -0.085 | 0.010 |
| Beta peak power (log) |  |  |  |  |  |  |
| EO (1) v EC (2) | 0.147 | 0.037 | <.001 | 0.075 | 0.219 | 0.197 |
| Age in years | <0.001 | 0.005 | .995 | -0.009 | 0.009 | <0.001 |
| Cardiometabolic burden | -0.189 | 0.049 | <.001 | -0.285 | -0.093 | 0.001 |
| APOE-ε4 | 0.147 | 0.079 | .062 | -0.007 | 0.302 | <0.001 |

\*EO=1 and EC=2; no E4 allele = 1, any E4 allele =2.

### 3.3. Associations Between Age, ACE-III Scores, Aperiodic Exponent and Alpha Peak Frequency Values

There was a significant negative correlation between age and alpha peak frequency (*r* = - 0.29, *p* = .007, BF_10_ = 5.05, depicted in Figure 2C). However, we did not detect a statistically significant association between age and the aperiodic exponent, with the Bayes factor indicating moderate support for the null hypothesis (*r* = -0.12, *p* = .287, BF_10_ = 0.24, Figure 2A). Please note that these are results of pairwise correlations which do not include other variables (such as cognitive performance) as covariates, following the approach in previous work (e.g., Voytek et al., 2015; Finley et al., 2022).

**Figure 2.**
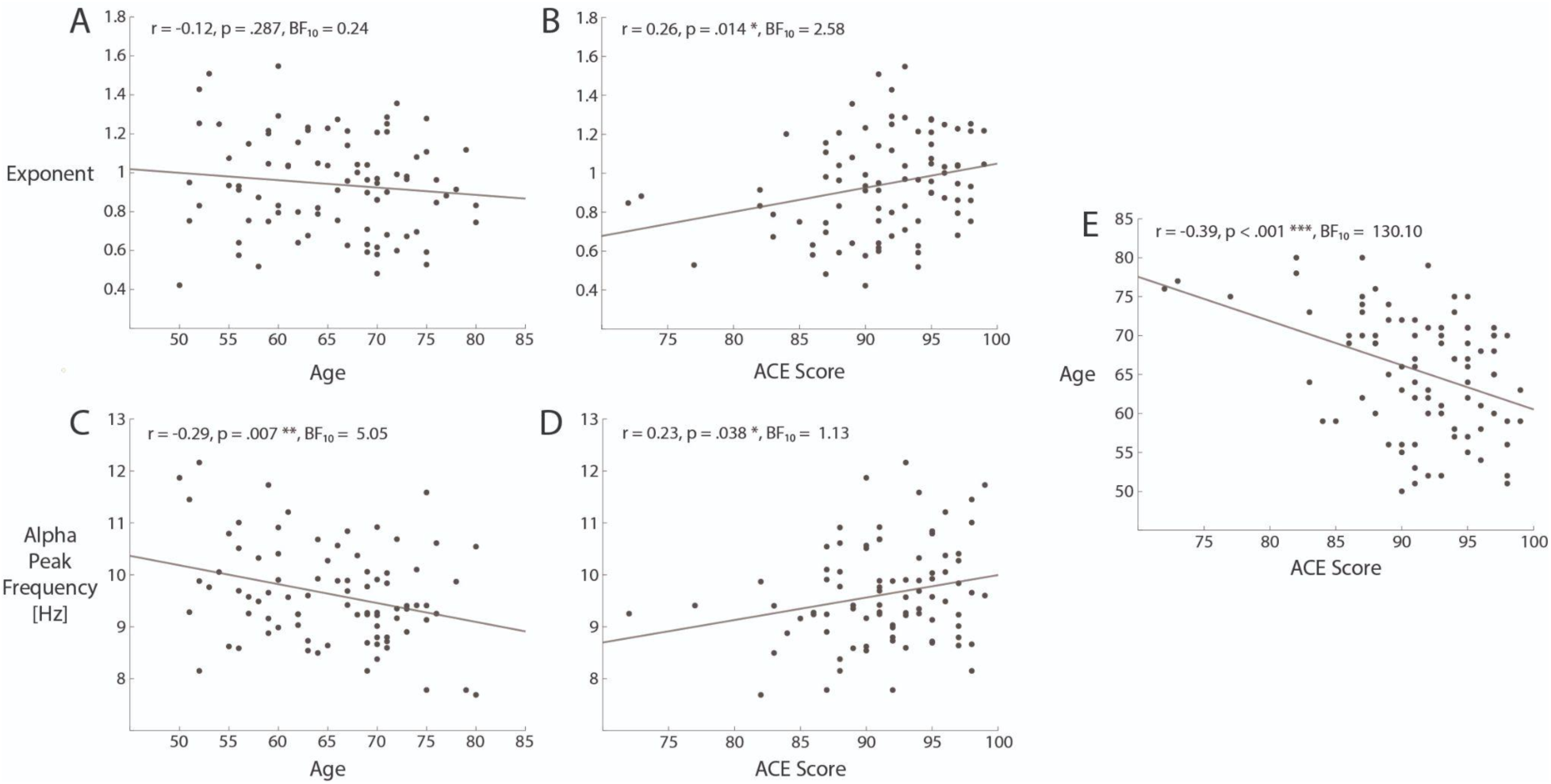
Associations between aperiodic exponent values, alpha peak frequency, age and ACE-III scores. Data points for EEG-derived measures were averaged across eyes open and eyes closed conditions within each participant. Gray lines depict lines of best fit from linear regressions including either ACE score, the exponent or alpha peak frequency as the sole predictor.

There were significant positive-going correlations between ACE-III total scores and the aperiodic exponent (*r* = 0.26, *p* = .014, BF_10_ = 2.58, Figure 2B) and alpha peak frequency (*r* = 0.28, *p* = .038, BF_10_ = 1.13, Figure 2D). Shapiro-Wilk tests indicated that data including ACE-III scores were not bivariate normal (p’s < .001), and so Spearman correlations were performed. The correlation with the exponent remained statistically significant (rho = 0.29, *p* = .007), however the correlation with alpha peak frequency was no longer significant (rho = 0.19, *p* = .078).

As expected, there were negative associations between age and ACE scores such that older adults showed worse cognitive performance, both when using Pearson (*r* = -0.39, *p* < .001, BF_10_ = 130.10) and Spearman (rho = -0.35, p = .001) correlations (Figure 2E).

To compare our results more directly to the primary analyses reported in Finley et al. (2022) we also measured aperiodic exponents at comparable electrodes to those in their analyses (Fz, F3 and F4). Exponents were not significantly correlated with age (*r* = -0.09, *p* = .425, BF_10_ = 0.18) but were positively associated with ACE scores (*r* = 0.25, *p* = .020, BF_10_ = 1.97). Please note that Finley et al. (2022) also report results from a whole-scalp composite ROI, similar to the ROI we used to measure the aperiodic exponent. Those results are very similar to the results from their primary analyses that only included frontal electrodes.

We additionally tested for correlations between aperiodic offsets, age and ACE scores in exploratory analyses, as lower offset values have been reported for older compared to younger adults (e.g., Donoghue et al., 2020a; Merkin et al., 2021). Offsets did not show statistically significant associations with age (*r* = -0.18, *p* = .090, BF_10_ = 0.56) or ACE scores (*r* = 0.20, *p* = .068, BF_10_ = 0.70).

To enable comparison with work using conventional band power analyses, we also tested for correlations between age, ACE scores, aperiodic exponent values, and power in the delta, theta, alpha and beta bands. We also performed exploratory analyses to test whether any measures from the parameterised power spectra (including alpha and beta peak power measures) were associated with age or ACE scores. Results are detailed in the Supplementary Material.

### 3.4. Associations Between Age, Cardiometabolic Burden, APOE-ε4 Carriage and EEG Measures

Table 2 displays the results from the five mixed effects models, with the outcomes of the exponent, alpha peak frequency, alpha peak power, beta peak frequency and beta peak power. Power spectra for the eyes open condition for different subgroups (including younger/older age and low/high ACE-III score subgroups) are plotted in Supplementary Figure S1. Participants with higher cardiometabolic burden had lower alpha peak frequencies (Figure 3A) as well as lower peak power measures in the beta band (Figure 3B). Participants with APOE-ε4 carriage also had lower beta peak frequencies (Figure 3C). Congruent with the correlation analyses, age was associated with alpha peak frequency but not the aperiodic exponent. Results for other measures were not statistically significant.

**Figure 3.**
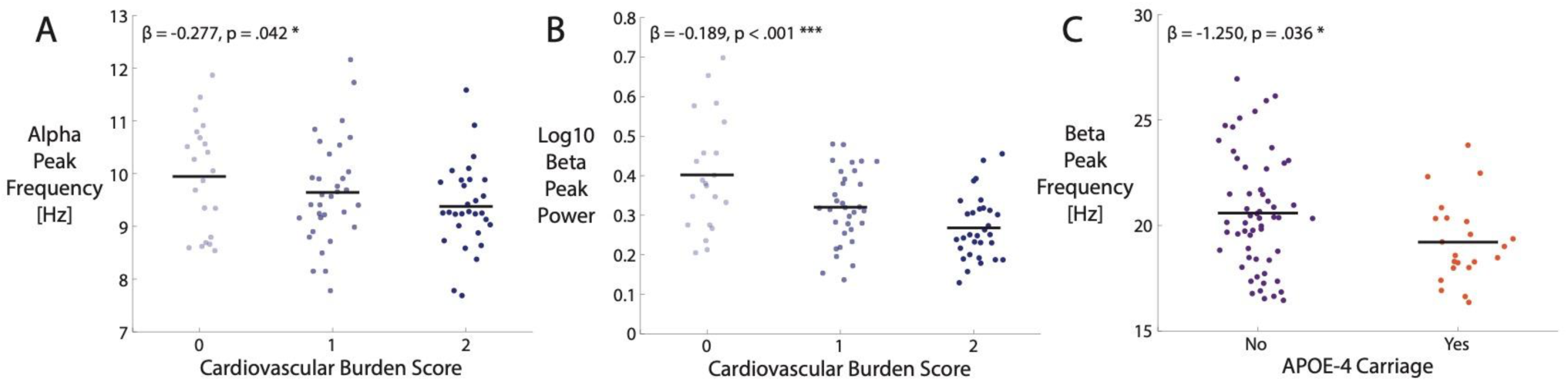
Statistically significant associations between EEG measures, cardiometabolic burden and APOE-ε4 carriage. A.) Alpha peak frequency values by cardiometabolic burden score. B.) Beta peak power (log-transformed) by cardiometabolic burden score. C.) Beta peak frequency by APOE-ε4 carriage. Black lines denote mean values for each group. Beta values and corresponding p-values are derived from the mixed effects modeling results in Table 2.

## Discussion

Our findings indicate that the slope of the aperiodic component of resting EEG power spectra is associated with measures of cognitive performance, but not chronological age, in adults aged between 50-80 years. Increased age was associated with slower alpha peak frequency but was not associated with flatter slopes of the aperiodic component. Dementia risk factors of APOE-ε4 carriage and poor cardiometabolic health displayed associations with periodic EEG measures in the alpha and beta bands. Taken together, our findings suggest that different risk factors for late-life cognitive decline and dementia exert partially overlapping effects on periodic and aperiodic resting EEG measures.

### 4.1. Associations Between Age, Cognitive Performance, Aperiodic Component Slope and Alpha Peak Frequency

Alpha peak frequencies were slower for older individuals, consistent with a large body of existing work (Klimesch, 1999; Grandy et al., 2013; Scally et al., 2018; Sghirripa et al., 2020; Merkin et al., 2021; Finley et al., 2022). By contrast, we found that the slope of the aperiodic component was associated with cognitive performance but not with chronological age. This finding is contrary to existing work comparing younger (e.g., 20-30 years) and middle-aged (e.g., 50-60 years) adults with relatively smaller sample sizes (e.g., Voytek et al., 2015; Dave et al., 2018), as well as a more recent study showing changes in aperiodic slope across 36-84 years of age in a larger (n=268) sample (Finley et al., 2022). However, our findings are congruent with existing work that also identified positive-going correlations between the steepness of the aperiodic slope and cognitive performance (Voytek et al., 2015; Ouyang et al., 2020).

There are a number of differences between our resting EEG study and previous work with respect to experimental task conditions (e.g., a working memory task in Voytek et al., 2015) and electrodes used to derive aperiodic slope measures (differing across studies), which complicates direct comparisons of results. However, our findings suggest that associations between cognitive ability and aperiodic slope measures may have been misidentified as effects of chronological age in previous work. This is because younger and older groups would be expected to differ with respect to both age and cognitive ability (due to age-related cognitive decline). The majority of previous work has not assessed relationships between cognitive task performance and age (Finley et al., 2022; Dave et al., 2018; Merkin et al., 2021; Scally et al., 2018; McNair et al., 2019; Koscieska et al., 2020). Those that did analyze task performance reported better performance in younger adult groups (Voytek et al., 2015; Leenders et al., 2018; Tran et al., 2020) congruent with our finding that older adults showed worse cognitive performance. Importantly, the slope of the aperiodic component was positively correlated with measures of processing speed in a large sample of younger adults (18-40 year-olds) in Ouyang et al. (2020), indicating an association with cognitive performance that is not specifically due to aging.

Another possibility is that aperiodic slope measures do substantially flatten over the course of early and middle-adulthood, but do not show such large magnitude changes during late life. Consequently, any associations with age (if they exist) may have been too small to detect in our sample of older adults. Notably, existing work has compared samples of young and middle aged adults (e.g., Voytek et al., 2015; Dave et al., 2018) or included younger participants than in our sample (e.g., Finley et al., 2022). The trajectory of the aperiodic slope may be nonlinear and partially distinct from age-related slowing of alpha peak frequency.

Our findings encourage a more nuanced interpretation of the neural noise hypothesis of aging, whereby the balance of excitatory and inhibitory neural activity (assumed to be indexed by the slope of the aperiodic activity, Hong and Rebec, 2012; Gao et al., 2017; Washke et al., 2017) is more specifically associated with cognitive ability, measured as performance on cognitive tasks. While cognitive ability typically declines with age, it can also vary substantially between individuals within a narrow age range. Other changes that occur with increased age, such as altered neurovascular coupling (Han et al., 2020) and differences in the morphology of brain tissue (e.g., Bedard et al., 2006; Bedard and Destexhe, 2009), could also plausibly contribute to individual differences in both neuronal noise and cognitive ability over the lifespan. Our findings highlight the importance of disentangling chronological age from other individual differences (such as cognitive ability) when testing and refining the neural noise hypothesis of aging.

### 4.2. Associations Between Resting EEG Power Measures, Cardiometabolic Burden and APOE-ε4 Carriage

We found that higher cardiometabolic burden was associated with lower alpha peak frequency and lower beta band peak power, and that APOE-ε4 carriage was associated with lower beta band peak frequency. Please note that, in our analyses, alpha and beta power values were corrected for the contribution of aperiodic EEG activity, in contrast to conventional frequency band power analyses. Our findings provide preliminary evidence that these modifiable and non-modifiable risk factors correlate with patterns of functional brain activity beyond concurrent effects of age. It is plausible that APOE-ε4 carriage and cardiometabolic burden increases neuroinflammation, tau hyperphosphorylation, Aβ aggregation and clearance, which are reflected in the EEG (e.g., Babiloni et al., 2006a, 2020). However, these mechanistic links were not directly explored here, and we caution that these findings should be replicated in an independent sample before strong conclusions can be drawn.

Whilst the mechanisms determining peak alpha frequency are yet to be fully elucidated, previous work suggests that decreases in peak alpha frequency may be a result of disrupted thalamo-cortical feedback loops, and/or deterioration of white matter and disconnection between neurons (Valdés-Hernández et al., 2010; Babiloni et al., 2020; Minami et al., 2020; Kumral et al., 2022). Notably, cardiometabolic risk factors (smoking, hypertension, pulse pressure, diabetes, hypercholesterolemia, body mass index and waist to hip ratio) have been associated with reduced brain volume, higher white matter hyperintensity volume, and poorer white matter microstructures in association with the thalamic pathways in large UK Biobank cohorts (n=9,722; Cox et al., 2019). This is one pathway by which structural changes may manifest as changes in measures of neuronal activity, highlighting the vulnerability of the brain to cardiometabolic factors across mid- to late-life.

We are hesitant to draw strong inferences about the functional significance of other changes in EEG spectral power profiles and how they relate to specific aspects of late life cognitive decline and dementia risk. This is because, in resting EEG studies, there is little control over cognitive or mental behavior during the EEG recording session. Differences in alpha or beta-band measures may have arisen due to changes in brain structure or function that are associated with APOE-ε4 carriage or increased cardiovascular burden, changes in cognitive activity during the resting EEG period (e.g., degrees of rumination, memory retrieval or future thinking) or a combination of these (discussed in Jach et al., 2020). Future work should complement resting EEG recordings with measures derived during cognitive tasks to assess whether these periodic measures also covary with specific measures of task performance.

### 4.3. Limitations

The findings of our study should be interpreted with the following caveats in mind. First of all, the FOOOF algorithm did not reliably identify peaks in the delta and theta bands in our dataset, meaning that we could not assess changes in periodic activity within these bands. Such peaks were also not clearly observable in the data of Merkin et al. (2021), who recruited participants of a similar age range, and could not be identified in the majority of participants in Finley et al. (2022). Peaks in the theta band may be best assessed in the context of working memory and decision-making tasks, which may evoke task-related theta-band activity that can be isolated and measured (e.g., Voytek et al., 2015; Cooper et al., 2015). These results also highlight the importance of accounting for aperiodic activity when measuring and interpreting EEG signals in the theta band (discussed in Finley et al., 2022).

In addition, there was a relatively low number of APOE-ε4 carriers in the recruited sample, meaning that we may not have had sufficient power to detect more subtle differences in resting EEG power measures. This is not surprising given that the estimated proportion of APOE-ε4 allele carriage is 13.7%, worldwide (Farrer et al., 1997). Much larger samples may be required to accurately characterize the profile of spectral changes that co-occur with APOE-ε4 carriage (e.g., Babilioni et al., 2006a).

We also note that we used general measures of cognitive performance assessed using the ACE-III. Consequently, it is unclear whether the aperiodic slope is related to performance in a subset of specific tasks (as implied in Voytek et al., 2015) or more general measures such as processing speed (as reported by Ouyang et al., 2020). Systematic investigations of covariation between aperiodic slopes and specific cognitive capacities may better pinpoint the functional significance of aperiodic activity and how it maps onto the neural noise hypothesis of aging.

Regarding our sample, we chose to include a broad spectrum of community dwelling older adults without dementia. This resulted in older adults across the spectrum of cognition and probably a mixture of neuropathologies that may contribute to future cognitive decline. The co-occurrence of such pathologies is present in many studies investigating older adults.

### 4.4. Conclusion

We report that, in a sample of 50-80 year-olds, the slope of the aperiodic component in resting EEG was associated with cognitive performance, but not with age. Our findings promote a more nuanced version of the neural noise hypothesis of aging and highlight ways to further develop and test this hypothesis. We also observed changes in periodic EEG activity with increasing cardiometabolic risk burden and APOE-ε4 carriage, suggesting that these dementia risk factors relate to observable changes in functional brain activity. Although these findings remain to be verified and further characterized in future work, they highlight brain mechanisms by which these risk factors relate to both structural and functional measures of brain activity.

## Acknowledgements

This study was funded by a UniSA Research Themes Investment Scheme Project Grant. AES was supported by an NHMRC-ARC Dementia Research Development Fellowship (GNT1097397). HADK was supported by a NHMRC Dementia Research Leadership Fellowship (GNT1135676). DF was supported by an Australian Research Council Discovery Early Career Research Award (DE220101508). Funding sources had no role in the design and conduct of the study, collection, management, analysis, and interpretation of the data, or the preparation of the manuscript.

