## Supplementary Material for "Resting EEG power spectra across middle to late life: Associations with age, cognition, APOE-ε4 carriage and cardiometabolic burden"

### Analyses Stratified by Subjects Above and Below the 92 ACE-III Cut-Off for MCI

Please note that no participants had a formal diagnosis of MCI in our sample, and that scores below the cut-off of 92 for the ACE-III are not by themselves sufficient to diagnose MCI.

**Supplementary table 1** *Participants scoring below the 92 ACE-II cut-off for MCI (n=42). Results from mixed models, with EEG indices as outcomes, and predictors including condition (eyes open/eyes closed), age, cardiometabolic burden and APOE-ε4 status. Statistically significant (<.05) p-values are shown in bold text.*

| | **beta** | **SE** | ***P*** | **95% CI** | | ***f²*** |
|---|---|---|---|---|---|---|
| **Exponent** | | | | | | |
| EO (1) v EC (2) | 0.163 | 0.036 | **<.001** | 0.092 | 0.233 | 0.475 |
| Age in years | -0.005 | 0.005 | .293 | -0.015 | 0.004 | <0.001 |
| Cardiometabolic burden | 0.059 | 0.054 | .274 | -0.047 | 0.164 | <0.001 |
| APOE-ε4 | -0.022 | 0.099 | .823 | -0.217 | 0.172 | 0.030 |
| **Alpha peak frequency** | | | | | | |
| EO (1) v EC (2) | -0.140 | 0.159 | .381 | -0.452 | 0.173 | 0.018 |
| Age in years | -0.020 | 0.016 | .219 | -0.051 | 0.012 | <0.001 |
| Cardiometabolic burden | -0.468 | 0.179 | **.009** | -0.819 | -0.118 | <0.001 |
| APOE-ε4 | -0.435 | 0.330 | .188 | -1.082 | 0.213 | 0.091 |
| **Alpha peak power (log)** | | | | | | |
| EO (1) v EC (2) | 0.623 | 0.097 | **<.001** | 0.433 | 0.813 | 0.983 |
| Age in years | -0.011 | 0.011 | .323 | -0.031 | 0.010 | <0.001 |
| Cardiometabolic burden | -0.187 | 0.118 | .114 | -0.419 | 0.045 | <0.001 |
| APOE-ε4 | 0.236 | 0.218 | .280 | -0.192 | 0.664 | 0.073 |
| **Beta peak frequency** | | | | | | |
| EO (1) v EC (2) | -1.128 | 0.339 | **.001** | -1.792 | -0.464 | 0.234 |

| | | | | | | |
|---|---|---|---|---|---|---|
| Age in years | 0.095 | 0.048 | **.048** | 0.001 | 0.189 | 0.258 |
| Cardiometabolic burden | 0.542 | 0.532 | .308 | -0.500 | 1.585 | <0.001 |
| APOE-ε4 | -2.112 | 0.978 | **.031** | -4.028 | -0.196 | 0.079 |
| **Beta peak power (log)** | | | | | | |
| EO (1) v EC (2) | 0.102 | 0.053 | .053 | -0.002 | 0.205 | 0.033 |
| Age in years | 0.004 | 0.007 | .573 | -0.009 | 0.017 | <0.001 |
| Cardiometabolic burden | -0.213 | 0.076 | **.005** | -0.362 | -0.065 | 0.001 |
| APOE-ε4 | 0.077 | 0.140 | .583 | -0.198 | 0.352 | <0.001 |

***EO=1 and EC=2; no E4 allele = 1, any E4 allele =2.***

**Supplementary table 2** *Participants scoring above 92 ACE-III cut-off for MCI (n=48). Results from mixed models, with EEG indices as outcomes, and predictors including condition (eyes open/eyes closed), age, cardiometabolic burden and APOE-ε4 status. Statistically significant (<.05) p-values are shown in bold text.*

| | beta | SE | *P* | 95% CI | | $f^2$ |
|---|---|---|---|---|---|---|
| **Exponent** | | | | | | |
| EO (1) v EC (2) | 0.144 | 0.047 | **.002** | 0.053 | 0.236 | 0.254 |
| Age in years | -0.002 | 0.005 | .712 | -0.012 | 0.008 | 0.003 |
| Cardiometabolic burden | -0.087 | 0.043 | **.043** | -0.171 | -0.003 | 0.005 |
| APOE-ε4 | 0.036 | 0.068 | .600 | -0.097 | 0.168 | 0.003 |
| **Alpha peak frequency** | | | | | | |
| EO (1) v EC (2) | -0.382 | 0.189 | **.043** | -0.753 | -0.011 | 0.105 |
| Age in years | -0.067 | 0.022 | **.002** | -0.111 | -0.024 | 0.005 |
| Cardiometabolic burden | 0.020 | 0.190 | .915 | -0.351 | -0.392 | 0.002 |

| | | | | | | |
|---|---|---|---|---|---|---|
| APOE-ε4 | -0.116 | 0.296 | .694 | -0.696 | 0.463 | 0.051 |
| **Alpha peak power (log)** | | | | | | |
| EO (1) v EC (2) | 0.724 | 0.088 | **<.001** | 0.552 | 0.895 | 1.792 |
| Age in years | 0.037 | 0.011 | **.001** | 0.016 | 0.059 | 0.004 |
| Cardiometabolic burden | -0.080 | 0.095 | .401 | -0.267 | 0.107 | 0.001 |
| APOE-ε4 | -0.232 | 0.146 | .111 | -0.512 | 0.054 | 0.018 |
| **Beta peak frequency** | | | | | | |
| EO (1) v EC (2) | -1.565 | 0.380 | **<.001** | -1.816 | -0.820 | 0.464 |
| Age in years | -0.018 | 0.056 | .749 | -0.019 | 0.124 | 0.002 |
| Cardiometabolic burden | 0.953 | 0.488 | .051 | -0.012 | 1.429 | 0.002 |
| APOE-ε4 | -0.527 | 0.731 | .780 | -1.959 | -0.904 | 0.064 |
| **Beta peak power (log)** | | | | | | |
| EO (1) v EC (2) | 0.201 | 0.049 | **<.001** | 0.105 | 0.297 | 0.448 |
| Age in years | -0.004 | 0.007 | .562 | -0.017 | 0.009 | 0.002 |
| Cardiometabolic burden | -0.150 | 0.059 | **.011** | -0.266 | -0.034 | 0.002 |
| APOE-ε4 | 0.176 | 0.089 | **.049** | 0.000 | 0.351 | 0.050 |

***EO=1 and EC=2; no E4 allele = 1, any E4 allele =2.***

### Analyses Using Conventional Resting EEG Band Power Measures

In addition to measures derived from parameterised power spectra, we also calculated conventional frequency band power measures as used in previous work. We calculated power across frequency bins within the delta (2-3 Hz), theta (4-7 Hz), alpha (7-14 Hz) and beta (15-30 Hz) bands separately. For alpha band power we used the same channels as for the parietooccipital region of interest (ROI) used to measure alpha peak frequency and power values in the parameterised power spectra. For all other measures, we used the same channels as the broad ROI used to measure the exponent in the parameterised spectra. As for the correlation analyses involving exponent and alpha peak frequency measures, data were first averaged across eyes open and eyes closed conditions within each participant. Results are summarized in Supplementary Table 3.

We tested for associations with age and Addenbrooke's Cognitive Examination (ACE) scores using Pearson correlations. Associations with age were not found for activity in the delta, theta, alpha or beta bands. Similarly, associations with ACE scores were not found for any frequency band.

To assess how the shape of the spectral profile of aperiodic activity relates to conventional band power measures we also tested for associations between each frequency band power measure and the aperiodic exponent. There were positive associations between exponent values and power in the delta and theta bands, and a negative association between exponent values and power in the beta band . There was no statistically significant association for power in the alpha band.

**Supplementary table 3** *Pearson's correlation analyses between conventional EEG power band measures and age, ACE Scores and aperiodic exponent values.*

| | **Age** | | | **ACE Scores** | | | **Aperiodic Exponent** | | |
|---|---|---|---|---|---|---|---|---|---|
| | ***r*** | **p** | **$BF_{10}$** | ***r*** | **p** | **$BF_{10}$** | ***r*** | **p** | **$BF_{10}$** |
| **Delta** | -0.02 | .858 | 0.14 | 0.07 | .524 | 0.16 | 0.51 | **< .001** | 31,267 |
| **Theta** | -0.05 | .670 | 0.15 | -0.01 | .906 | 0.14 | 0.42 | **< .001** | 413.62 |
| **Alpha** | 0.02 | .831 | 0.14 | -0.02 | .863 | 0.14 | 0.16 | .137 | 0.40 |
| **Beta** | -0.17 | .113 | 0.46 | -0.12 | .292 | 0.23 | -0.34 | **.001** | 22.92 |

### Correlations Between Parameterised Power Spectra Measures, Age and ACE Scores

We additionally tested for associations between each measure derived from the parameterised power spectra, age and ACE scores in a set of exploratory analyses. We used Pearson correlations for all analyses reported here. Please note that results for the aperiodic exponent, aperiodic offset and alpha peak frequency are also reported in the paper.

Results are summarized in Supplementary Table 2. There were no statistically significant associations with either age or ACE scores for alpha peak power, beta peak power or beta peak frequency measures.

**Supplementary table 4** *Pearson's correlations between measures from parameterised power spectra, age and ACE scores.*

| | **Age** | | | **ACE Scores** | | |
|---|---|---|---|---|---|---|
| | ***r*** | **p** | **$BF_{10}$** | ***r*** | **p** | **$BF_{10}$** |
| **Aperiodic Exponent** | -0.12 | .287 | 0.24 | 0.26 | **.014** | 2.58 |
| **Aperiodic Offset** | -0.18 | .090 | 0.56 | 0.20 | .068 | 0.70 |
| **Alpha Peak Power** | 0.14 | .200 | 0.31 | 0.01 | .954 | 0.14 |
| **Beta Peak Power** | -0.06 | .603 | 0.16 | 0.08 | .491 | 0.17 |
| **Alpha Peak Frequency** | -0.29 | **.007** | 5.05 | 0.28 | **.038** | 1.13 |
| **Beta Peak Frequency** | 0.15 | .170 | 0.34 | -0.07 | .509 | 0.17 |

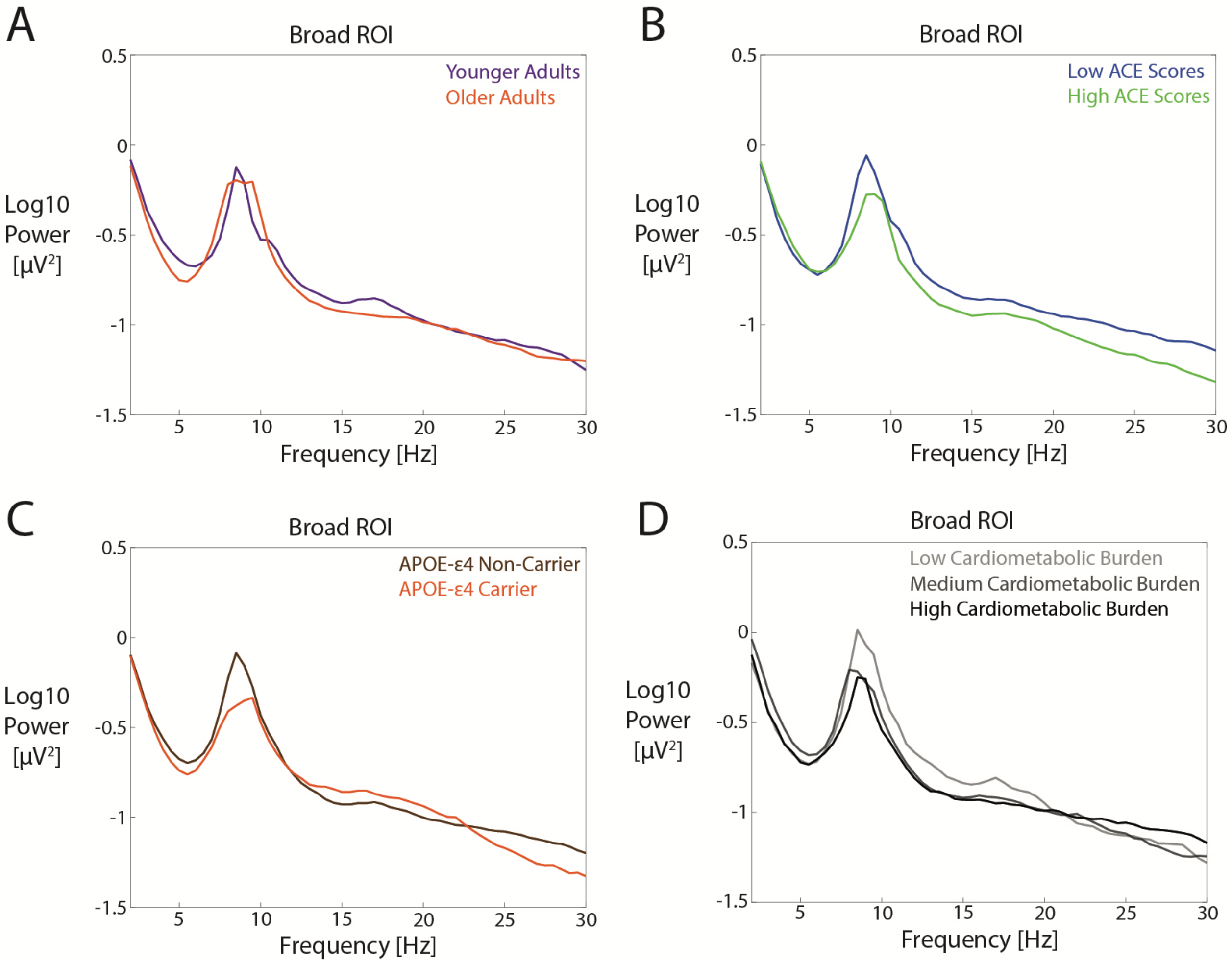


**Supplementary Figure S1.** Fourier power spectra for the eyes open condition plotted by subgroup. A.) Younger and older adults, with subgroups defined by a median split. B.) Relatively low and high ACE-III scores, defined by a median split. C.) APOE-ε4 carriers and non-carriers. D.) Low, medium and high cardiometabolic burden groups, defined by the numbers of recorded risk factors as defined in the methods. Data were averaged across all electrodes comprising the broad ROI (Fz, F3, F4, Cz, C3, C4, Pz, P3, P4, PO3 and PO4).
